# Complexity as a Potential Neurophysiological Correlate of Awe

**DOI:** 10.1101/2025.11.20.689403

**Authors:** Joseph CC Chen, Gabriella Mace, Avery Ostrand, Christian Valtierra, Sydney Griffith, Richard Campusano, Andrew Li, Erin Vinson, Jonas TT Schlomberg, Maya Eshel, John M. Suntay, Syed Rahim, Roger Anguera-Singla, Bijurika Nandi, Isabella Hartley, Daniel A. Brown, Nicole C. Swann, Luca Mazzucato, Xin Hu, Dan Zhang, Christopher Timmermann, Maryam Bijanzadeh, Reza Abbasi-Asl, Robin L Carhart-Harris, Theodore Zanto, David A Ziegler, Adam Gazzaley, Lorenzo Pasquini

## Abstract

Awe is a positive emotion often accompanied by sensations of vastness and unity, with known benefits for well-being and social behavior. However, its neural underpinnings remain poorly understood. We recorded electroencephalography (EEG) and autonomic physiology in 23 healthy older participants while they watched a nature-based audiovisual film and subjectively rated awe events. Awe was the predominant emotion reported – though other positive emotions (e.g., joy) were also highly rated. Awe events were associated with decreased skin conductance level (SCL), and decreased EEG alpha and theta spectral power – physiological changes associated with low arousal and positively valenced emotional states. Interestingly, awe events exhibited increased Lempel Ziv Complexity (LZC) – indicating heightened neural signal entropy and increased richness of the conscious experience. LZC was also positively associated with the intensity of the awe ratings and negatively associated with SCL. Three additional datasets with separate independent induction methods (video clips and pharmacological induction via N,N-dimethyltryptamine) also showed positive occipital LZC associations with awe – suggesting some generalizability of LZC as a neurophysiological marker of awe. These results suggest that awe evokes neurophysiological states linked to the subjective affective experience.

**Impact Statement:** Awe, an emotion of increasing interest, has been studied with fMRI and peripheral physiology – but few studies have used electroencephalography (EEG). In this EEG study, we utilize a movie-watching paradigm to explore potential physiological correlates of awe. The results present EEG-based complexity increases as a potential correlate of awe, though showing limited generalization to independent datasets and limited uniqueness compared to joy, another positive emotion.

## 1. Introduction

Awe is a positive emotion, characterized by feelings of wonder and unity, often experienced when encountering something vast and complex, such as a breathtaking vista, engaging with complex art or music, or witnessing acts of extraordinary kindness or courage (Keltner & Haidt, 2003; Monroy & Keltner, 2022; Shiota, 2021; Shiota et al., 2007). The experience of awe can induce a feeling of being part of something larger than ourselves, prompt a need for accommodation of prior beliefs (Keltner & Haidt, 2003; Monroy & Keltner, 2022; Shiota, 2021; Shiota et al., 2007), which can help move away from a self-centered perspective (Chirico & Yaden, 2018; Monroy & Keltner, 2022). This cognitive shift can lead to increased empathy and a stronger sense of connection with others and the world around, which can be harvested to promote well-being as well as prosocial behavior (Guan et al., 2019; Jiang & Sedikides, 2022; Piff et al., 2015; Rudd et al., 2012; Stellar et al., 2017; Telle & Pfister, 2016; Van Cappellen & Saroglou, 2012).

Indeed, recent interventional studies leveraging the experience of awe, typically in the context of nature exposure, have shown that awe promotes improvements in well-being across various healthy and clinical populations (Anderson et al., 2018; Leavell et al., 2019; Lopes et al., 2020; Monroy et al., 2025; Monroy & Keltner, 2022; Sturm et al., 2020). For example, awe experienced during water rafting has shown to reduce stress, as well as improve overall well-being in military veterans, youth from underserved communities, and patients living with long COVID (Anderson et al., 2018; Leavell et al., 2019; Lopes et al., 2020; Monroy et al., 2025). When looking across the age span, awe-inspired interventions have been shown to increase well-being in both undergraduate students and older adults (Bai et al., 2021; Bratman et al., 2021; Monroy & Keltner, 2022; Sturm et al., 2020). Together, this work highlights awe as a scalable, non-pharmacological tool with wide-reaching benefits across the lifespan.

In the context of research, awe is typically elicited by either exposing study participants to nature (Hartig et al., 2014; Kuo, 2015) or by leveraging naturalistic paradigms (Chirico et al., 2017; Gross & Levenson, 1995; Ng et al., 2023; Pasquini et al., 2023; Saarimäki, 2021; Shiota et al., 2007), where participants are instructed to attend to awe-inducing audiovisual stimuli in a controlled laboratory setting (van Elk et al., 2019). Recently, awe has also been induced through the administration of psychedelic substances, such as psilocybin and N,N-dimethyltryptamine (DMT) (Timmermann et al., 2023), a class of serotonergic agonists producing rapid and durable improvements in mental health (Bogenschutz et al., 2022; R. Carhart-Harris et al., 2021; Falchi-Carvalho et al., 2025; Goodwin et al., 2022; Griffiths et al., 2016; Raison et al., 2023), believed to be partially mediated through feelings of awe commonly experienced under the effects of psychedelics (Goldy et al., 2024; Hendricks, 2018; Lebedev et al., 2015; Millière, 2017; Yaden et al., 2024).

One mechanism through which awe has been proposed to promote well-being and prosocial behavior is by inducing shifts in neurophysiology (Keltner & Haidt, 2003; Monroy & Keltner, 2022; Shiota, 2021; Shiota et al., 2007). Studies leveraging autonomic recordings have consistently shown reduced sympathetic outflow and increased parasympathetic activity in the context of awe, as indexed by reductions in skin conductance, heart rate, and respiration rate, as well as increases in heart rate variability (Pasquini et al., 2023; Shiota et al., 2011). From a central nervous system perspective, studies using functional magnetic resonance imaging (fMRI) have revealed that awe deactivates regions spanning the medial parietal and medial frontal cortices, which belong to a brain network supporting self-referential functions referred to as the default-mode network (Buckner & DiNicola, 2019; Takano & Nomura, 2022; van Elk et al., 2019).

Yet, little is known about the neurophysiological correlates of awe, despite electroencephalography (EEG) being a more direct measure of neuronal activity than fMRI or autonomic physiology (Mizuno-Matsumoto et al., 2020; Mulert, 2013), as well as being better posed to measure rapid changes in brain activity that may capture dynamic emotional states, including awe (Gross, 2015; Monroy & Keltner, 2022). Oscillatory EEG changes can be observed in affective contexts – for example, alpha desynchronization was observed during anxiety elicited by angry facial presentations (Knyazev et al., 2008), greater feelings of humiliation (Otten & Jonas, 2014, p. 201), and in a review of studies on picture processing suggesting that alpha desynchronization indexes emotional processing (Codispoti et al., 2023). Conversely, theta and alpha synchronization has been observed in high arousal stimuli (Aftanas et al., 2002). In parallel, emerging EEG approaches such as Lempel Ziv Complexity (LZC) quantify the entropy or uncertainty in time series data and are increasingly used in neuroscience research. LZC has been shown to differ across varying states of sleep, wakefulness, attention tasks, memory tasks (Höhn et al., 2024), and during psychedelic states (Schartner, Carhart-Harris, et al., 2017; Schartner, Pigorini, et al., 2017). While LZC is hypothesized to capture heightened information processing and the richness of the conscious experience (R. L. Carhart-Harris, 2018; R. L. Carhart-Harris et al., 2014), no studies, to our knowledge, have investigated the effect of emotions on LZC.

Here, we capitalized on EEG, autonomic physiology, and self-report data acquired during naturalistic audiovisual stimulation in a sample of 23 healthy volunteers, revealing reduced EEG alpha and theta power, as well as increased LZC. To explore the generalizability of awe-induced neurophysiological changes, we included three external EEG datasets, two leveraging audiovisual stimuli (X. Hu et al., 2017) and a third administering the psychedelic DMT (Timmermann et al., 2019), to explore the convergence of awe-induced neurophysiological changes across samples, experimental settings, and modes of awe induction. Finally, we capitalize on the self-report and autonomic data simultaneously acquired with EEG in the primary dataset, to investigate the link between neurophysiological changes, reduced sympathetic activity, and strength of experienced awe.

## 2. Methods

The study protocol was approved by the University of California San Francisco Institutional Review Board (22-37461) and was conducted in accordance with the Declaration of Helsinki. All participants provide informed consent. The study was registered as part of a larger ongoing clinical trial (https://clinicaltrials.gov/study/NCT05645835). The code used to analyze is uploaded in https://github.com/josephccchen/dynamo.

### 2.1. Participants

Twenty-three cognitively intact older adults were recruited (ages ranging from 60 – 85) with their details, demographics, and cognitive scores described in Supplementary Table 1**Error! Reference source not found.**. Participants were recruited from a registry of older adult control volunteers, who had been assessed with neuropsychological tests of executive and memory function via a previously described remote characterization module (Arioli et al., 2022) at the University of California San Francisco Neuroscape Center. The neuropsychological evaluation administered working memory and verbal learning tests as in the CVLT-II (Delis et al., 1987), processing speed (WAIS-R) (Wechsler, 1997), visual-motor sequencing (DKEFS Trail-Making A and B), semantic fluency and phonemic fluency (DKEFS) (Delis et al., 2001). Furthermore, as research measures tracking self-referential functions and social well-being, the UCLA Loneliness scale (D. W. Russell, 1996) and the Reflection-Rumination Questionnaire (RRQ) (Trapnell & Campbell, 1999) were administered to participants.

### 2.2. Primary Awe Paradigm

Participants were seated in a quiet, dark room in front of a 27-inch monitor and dual speakers. A 29-minute nature video, *The Nature Journey,* directed by Louis Schwartzberg and with music by East Forest was played to participants twice. The first “passive” viewing was played uninterrupted from beginning to end with participants rating *The Nature Journey* on ten emotions at the end using a 10-point visual analog scale. Participants used the left and right arrow keys on a keyboard to rate affection, amusement, joy, pride, surprise, awe, anger, anxiety, boredom, disgust, fear, sadness, arousal, and valence.

During the second viewing of *The Nature Journey*, participants were instructed to pause every time they remember experiencing awe (Figure 1A) using the Spacebar key on their keyboards. Participants then rated the intensity of awe on a 10-point visual analog scale using the right and left arrow keys, then resume by pressing the Return key. The timestamps of these individual awe ratings were automatically stored in a text file and then projected onto the first passive viewing to determine the EEG epochs corresponding to awe events experienced by the participant during the first viewing (Figure 1B).

**Figure 1.**
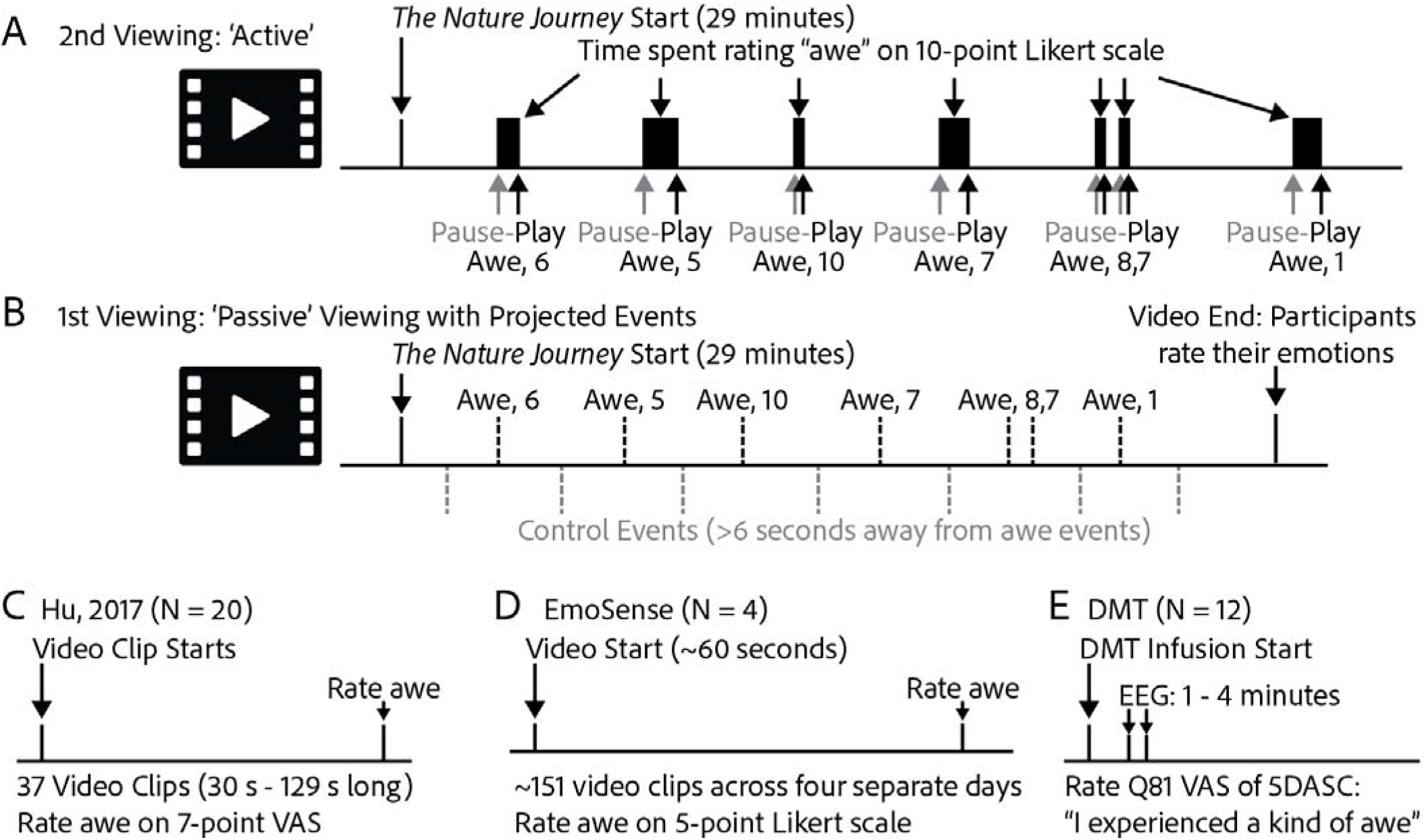
Study Design. (A) After participants passively watched a 29-minute nature-inspired audiovisual film – The Nature Journey – while their brain activity was recorded with EEG and autonomic recordings, participants watched The Nature Journey a second time and self-reported when they remembered experiencing awe and at what intensity (1-10 visual analog scale). (B) These awe events were projected back into the passive viewing and random control events were identified for each participant, as timepoints at least 6 seconds away from awe events and other control events. To assess generalizability of awe, three additional datasets were consulted: (C) The first study measured EEG in twenty healthy Chinese volunteers while participants watched 37 video clips and rated ten positive emotions (Hu, 2017). (D) The second study measured EEG in four healthy adults while participants watched ∼151 60-second video clips and rated awe on a 5-point Likert scale. (E) The third study acquired EEG during N,N-dimethyltryptamine (DMT) infusion. EEG data from the first 1-4 mins after the intravenous administration of the psychedelic DMT was used to assess the neurophysiological correlates of awe induced through a pharmacological approach instead of an audiovisual stimulus. Participants in this study filled out the 5DASC questionnaire, of which one question (Q81) asked participants whether they agreed or disagreed with the statement “I experienced a kind of awe” on a visual analogue scale (VAS).

The task was designed in Unity (https://unity.com/), which was programmed to store the self-report data. The Unity task was programmed to send photodiode triggers to both the BIOPAC MP160 system (Biopac Systems Inc., Goleta, CA, https://www.biopac.com/) and Biosemi ActiveTwo system (Biosemi BV, Amsterdam, Netherlands, https://www.biosemi.com/) systems for EEG and electrodermal data acquisition (further details below). The corresponding times at which participants started watching *The Nature Journey*, as well as when participants paused to rate awe and subsequently resumed were captured by the photodiodes.

### 2.3. Awe and Control Events Preprocessing

The control events were determined by pseudorandomly sampling events that fulfilled the following conditions: (i) >6 seconds away from any awe event, (ii) >6 seconds away from muscle artefacts (see section below for EEG muscle artefacts), (iii) >6 seconds away from either the beginning or the end of *The Nature Journey*, or lastly (iv) >6 seconds away from any other control event. Control events were reproducibly selected by using a seed and repeatedly selected until no more valid control events could be selected (Figure 1).

### 2.4. Characterization of Joy in *The Nature Journey*

In efforts to identify whether the neurophysiological markers were specific to awe, a secondary study was performed. Participants were asked to rewatch *The Nature Journey* online twice. The first time was a passive rewatching, while during the second rewatching, participants were asked to pause to rate whenever they remembered feeling the emotion of joy. The task was reprogrammed in Unity to show *The Nature Journey* online so participants could watch from home. Twenty participants of the original twenty-three participated.

### 2.5. Influence of Visual Features

To assess for changes in visual features across awe and control events, continuous estimates of image luminosity and contrast were estimated for every frame of *The Nature Journey* by loading the video file into MATLAB (MathWorks, Boston, MA) and calculating the mean intensity (luminosity) and the standard deviation of the intensity of every frame (contrast). The luminosity and contrast for every awe and control event were then estimated by taking the average luminosity and contrast of all frames within 3 seconds of each awe and control event. Event-averaged levels of luminosity and contrast were statistically compared across awe and control events using type 3 analysis of variance of mixed effect models (*p*<0.05). Furthermore, to identify whether *The Nature Journey* characteristics (luminosity and contrast) affected EEG metrics (LZC, theta power, and alpha power), linear mixed-effect models were performed with random intercepts. This was performed by a two-way mixed effect model where the EEG metrics were predicted by the event type (awe or control) and video characteristic (luminosity or contrast; Supplementary Figure S1).

### 2.6. Independent Characterization of The Nature Journey

To assess whether *The Nature Journey* induced the experience of constructs relevant to the experience of awe, an additional experiment was performed. Six volunteers (mean ± standard deviation age = 25.3 ± 3.1, 4 males, 2 females), rated every scene of *The Nature Journey* according to six constructs: vastness, unity, connection to nature, aesthetic beauty, extraordinary, and complexity (Supplementary Figure S2). These ratings were used to establish an averaged time-series of each of these constructs and compare their intensities across awe and control events (determined by the study participants of the present report) using linear mixed effect models.

### 2.7. Neurophysiological Data Collection, Preprocessing, and Analyses

Electroencephalography was acquired from participants using 64-channel headcaps and Biosemi ActiveTwo amplifiers (Biosemi BV, Amsterdam, Netherlands). EEG was acquired at 2048 Hz with 24-bit resolution and were band-pass filtered between 0.01 Hz and 100 Hz during data acquisition.

Preprocessing steps of the EEG data were performed using FieldTrip (Oostenveld et al., 2011) (https://www.fieldtriptoolbox.org/) including within-channel demeaning, high-pass filtering at 0.5 Hz, semi-automated muscle artefact rejection, bad channel rejection, ocular independent components analysis rejection (with an average of 2.2 components removed per 29 minute recording), bad channel interpolation, average referencing, and down sampling to 500 Hz. Time frequency analysis was performed with a single Hanning taper, characterizing frequencies from 2 – 30 Hz in 0.1 Hz increments, and times from -3 to +3 s from the awe/control events in 0.02 s increments to avoid edge artifacts. The windows were set at three cycles of frequency of interest, though zero padded to the next power of two sample points. Only spectral power values from -3 to +3 s from the awe/joy/control events were retained for further analysis. The averaged time frequency representation of awe, joy, and control events were obtained for each participant, averaged over all channels, then statistically compared via a paired *t*-test. Visual inspection of the final statistical time frequency representation of awe - control revealed significantly decreased theta (5 – 8 Hz) and alpha (8 – 13 Hz) power from -2.8 s to 0.6 s from the awe/control events. The theta and alpha power values between -2.8 s to 0.6 s from the awe/control events were averaged across all awe and control events for each participant, then a paired *t*-test was performed between the 23 individual pairs of awe and control topographies. Non-parametric cluster-based permutation tests were performed to identify significant topographical clusters between the awe and control topographies with 5,000 random permutations. Analogous approaches were used when comparing the joy and awe topographies.

LZC analysis was performed on the 6 second epochs in a similar time-frequency manner where sliding windows of 6 seconds estimated LZC from -3 to +3 seconds in 0.02 s increments relative to each awe/control event. The EEG time series was binarized by the median value and the Lempel Ziv algorithm (Lempel & Ziv, 1976) was applied to the binary symbols to find the longest unique string of symbols. The normalized LZC was then calculated by dividing the longest string by the total length of the sequence (J. Hu et al., 2006). The normalized LZC (henceforth referred to as LZC) was then averaged across all awe, joy, and control events for each participant and statistically compared using a paired *t*-test (*p*<0.05, cluster-corrected). Non-parametric cluster-based permutation tests were performed to identify significant topographical clusters between the awe and control LZC topographies with 5,000 random permutations.

EEG channels belonging to significant topographical clusters were identified for the theta power, alpha power, and LZC. Subsequently, the average theta power, alpha power, and LZC value from those channels were derived. To identify whether the neurophysiological measures predicted the intensity of awe ratings, three separate linear mixed effect models (*p*<0.05) were performed to assess whether alpha power, theta power, or LZC values predict awe intensity values using *lme4* (Bates et al., 2015) in R (https://www.r-project.org/) to determine the interrelation between theta power, alpha power, and LZC values. A causal mediation analysis was performed using the ‘mediation’ R package (Tingley et al., 2014) to further assess whether the link between LZC increases and decreases in theta power was partially mediated by changes in the faster alpha power (*p*<0.05).

Furthermore, to assess the extent by which EEG metrics were influenced by visual features (e.g., contrast and luminosity), analyses controlling for luminosity and contrast were included in the linear mixed effect model and reported in Supplementary Figure S1.

### 2.8. MRI Data Collection and EEG Source Reconstruction

T1-MPRAGE images of participants’ brains were acquired with a 3T Prisma Fit (Siemens, Erlangen, Germany) with a 64-channel head coil at a separate visit for all but two participants. The pulse sequence parameters included TR = 2300 ms, TE = 2.96 ms, TI = 900 ms, flip angle = 9°, isotropic voxel sizes of 1.0 mm, and with GRAPPA acceleration factor 2. A field-of-view of 160 mm x 240 mm was acquired for 256 axial slices. For the two participants who did not have an MRI due to participant scheduling conflicts, standard MNI templates were used.

Source-based analyses were performed in FieldTrip along with its integration with SPM. The MRI was aligned and segmented into five tissue types: gray, white, cerebrospinal fluid, skull, and scalp. A finite element head model was created using the FieldTrip-SimBio pipeline (Vorwerk et al., 2018) whereby the conductivities of the five tissues types were assumed to be: 1.79, 0.33, 0.43, 0.01, 0.14 S/m respectively. Standard electrode positions were projected onto the scalp surface, and a template 10 mm grid of sources was overlaid on each individual participant. A linearly constrained minimum variance beamformer was used to reconstruct the sources and ascertain the EEG time series in each source location. Theta power, alpha power, and LZC was then calculated for each source location using the same parameters as described previously in sensor space and normalized into standard space. The averaged awe and control sources were statistically compared via a paired *t*-test. To ascertain which networks were implicated in awe, the statistical maps were interpolated to obtain t-statistic estimates according to the Schaefer-100 cortical atlas (Schaefer et al., 2018). For visual purposes, the awe – control statistical map was smoothed and interpolated on a standard cortical surface using BrainNet (https://www.nitrc.org/projects/bnv) (Xia et al., 2013).

### 2.9. Generalizability of Awe Neurophysiology Across Paradigms

Four additional datasets shared by collaborators were analyzed to assess the generalizability of neurophysiological findings. Three of the datasets directly include awe ratings, whereas a fourth dataset assessed goosebumps – a related but distinct phenomena on one participant, which was included as supplementary analyses. The aim of including additional awe paradigms was to investigate whether independent experiments on different samples would yield similar neurophysiological correlates of awe as in the primary study. To that end, the additional datasets were used to assess whether correlation of awe and EEG metrics yielded similar topographies.

#### Dataset 1 (Hu et al., 2017, N = 20)

The first dataset (X. Hu et al., 2017) involved twenty Chinese participants each being shown 37 film clips of positive emotions which were on average 72 s long. After each video clip, participants rated the strength of 10 positive emotions using 7-point continuous Likert scales. EEG with 32-channels were recorded from participants during the audiovisual presentation via a portable wireless EEG amplifier (NeuSen.W32, Neuracle, China). The EEG data from the last 30 seconds of each film were sliced into 5 s epochs with 50% overlap. For each epoch and each channel, theta power, alpha power, and LZC values were calculated.

#### Dataset 2 (Emotion Rating Study (ERS), N = 4)

The second dataset involved yet to be published data, collected by authors RC, BN, IH, MB, RA. EEG was recorded while four participants watched, on average, ∼151 60-second video clips each across four separate days. The video clips were assembled from a validated normative library of short emotional video clips each with normative affect ratings (Cowen & Keltner, 2017). Using 18 positive emotions out of 27 distinct emotion categories, the most representative clips were selected for each emotion by taking the top 20 highest normative-rated exemplars. Because a video clip could rank highly for more than one emotion, overlaps were resolved to enforce mutual exclusivity: any clip appearing in multiple emotions’ lists was retained only for the emotion in which its rating was highest, removed from the others, and each emotion’s list was backfilled with the next-highest ranked clip, iterating until every emotion had a disjoint set of slips and each clip assigned to a single emotion. The video clips selected for each emotion were then concatenated end-to-end into an emotion stimulus, yielding the ∼1-minute trial videos presented during the experiment with each clip composition and ordering of each trial fixed in advance. The ratings of awe were performed on a 5-point Likert scale for each video clip. The EEG recorded included 128-channels with the Biosemi ActiveTwo system.

#### Dataset 3 (“DMT”, N = 12)

The third dataset involved previously published data (Timmermann et al., 2019) where twelve participants were administered intravenous saline placebo and intravenous DMT, a serotonergic-2A agonist psychedelic, on two separate dosing visits separated by a week. DMT is a fast-acting psychedelic which reportedly induces intense feelings of awe (Goldy et al., 2024; Hendricks, 2018; Yaden et al., 2024). EEG data was acquired with 32-channels during the saline and DMT infusions via BrainProducts EasyCapMR32 (BrainProducts GmbH, Gilching, Germany). EEG data from the 1 – 4 minutes post-infusion time period were used to calculate theta power, alpha power, and LZC values. Participants also completed the 5-dimensional altered states of consciousness (5DASC) questionnaire (Dittrich, 1998), which contains 94 statements that are rated on a visual analogue scale where the left edge denoted “No, not more than usually”, and the right edge denoted “Yes, much more than usually”. Question 81 of the 5DASC asks “I experienced a kind of awe” and this rating was used to correlate with the EEG metrics.

#### Analysis

Because these paradigms and measures were collected in different contexts and different systems, standardized beta coefficients were analyzed from linear mixed effect models by z-scoring both EEG metrics and awe ratings within each subject prior to submitting to either a linear mixed effect model with random intercepts (for the current study and Datasets 1 and 2), or a simple linear model (for Dataset 3, DMT). The random intercepts were utilized to account for the repeated sampling from an individual participant. This analysis was performed per-channel and the topography of the standardized beta estimates was plotted.

To assess spatial similarity between each study’s topography for alpha power, theta power, and LZC, the standardized beta estimate at each common channel was pair-wise Pearson correlated. To assess whether the correlation metric surpassed chance, permutation-based tests shuffling the channel values with 10,000 iterations were performed to establish a null distribution, and the proportion of times the correlation metric was larger in magnitude than the null distribution was used to assess significance. Permutation-based significance testing was performed instead of using the analytical *p*-value due to the spatial autocorrelation of values across the EEG topography.

#### Dataset 4 (“Goosebumps”, N = 1)

The fourth dataset involved pilot data collected by authors DB, LM, and NS. Informed consent was obtained from one participant (34 years, male). A 40-minute audiovisual film designed to induce goosebumps was presented to this participant three times. The participant was instructed to press the spacebar whenever the participant experienced goosebumps. The audiovisual clips included short films, timelapses of nature, and ensemble music performances. EEG with 64-channels were recorded alongside the audiovisual stimuli. As previously described, control events were randomly sampled if these were at least 6 s from goosebump events, muscle artefacts, or other control events – until the number of control events and goosebump events matched. Theta power, alpha power, and LZC values were determined from goosebump and control events, averaged within participants, then statistically compared via paired *t*-test. The results are described in Supplementary Figure S3. The aim of this dataset was to assess the EEG correlates of goosebumps – an awe-related phenomena.

### 2.10. Electrodermal Data Collection, Preprocessing, and Analyses

Electrodermal activity was acquired using a BIOPAC MP160 system (Biopac Systems Inc., Goleta, CA) and EDA100C amplifier module. Isotonic electrolyte gel was applied and electrodermal activity electrodes were placed on ventral aspect of the first two digits of the participant’s left hand.

The electrodermal activity data was visually inspected for signs of expected fluctuation (i.e., to filter out non-responders to the electrodermal activity signal). Due to visual artifacts in the data, only data for 12 participants were retained for subsequent analyses. The 30-minute electrodermal activity signal was then z-normalized (by subtracting the mean and dividing by the standard deviation). From the electrodermal signal, the skin conductance level (SCL, in *z* units) within -3 to +3 s of each awe and control event was averaged to obtain one value per event.

Average SCL values were compared between awe and control events through linear mixed effect models with random intercepts per participant (*p*<0.05). SCL values from awe events were used to predict EEG metrics. These analyses revealed a significant association between SCL and LZC. Finally, we examined if the relationships between LZC and SCL values were influenced by luminosity. To this end, we performed a two-way interaction mixed effect model (LZC ∼ SCL + Luminosity).

### 2.11. Supplementary Analysis Examining Awe and LZC as Trait or Transient Metrics

To assess whether LZC under awe reflected a trait characteristic rather than a short-lived, transient, physiological state, we correlated the awe and LZC measures against questionnaires tracking self-referential functions and social well-being. Partial correlation was performed by removing the variance attributable to age, sex, years of education, and ethnicity prior to performing a correlation of residual against the measure of interest. The UCLA Loneliness scale (D. W. Russell, 1996) and Reflection-Rumination Questionnaire (RRQ) (Trapnell & Campbell, 1999) measures were partially correlated against the mean awe and mean LZC measures averaged across all awe events.

## 3. Results

### 3.1. Characterization of Awe in *The Nature Journey*

Participants of the present study rated awe, on average, as the most predominant emotion elicited by *The Nature Journey* compared to all other emotions (Figure 2A). The second-highest rated emotion after awe was joy – which was why a secondary experiment asked participants to rate joy. There were significantly larger numbers of identified control events compared to participant reported awe and joy events, though the number of awe and joy events did not significantly differ (Figure 2B). The distribution of awe and joy events appear to be similarly distributed across *The Nature Journey*, with control events expectedly occurring in times less associated with awe and joy (Figure 2C). In the additional experiment where participants rated *The Nature Journey* at home for joy events, the average rating of awe ratings remained the most intense (Figure 2D). Independent characterizations of scenes showed significantly higher ratings of complexity, extraordinary, and unity for awe and joy when compared to control events, but lower ratings of vastness (Figure 2E). The full, independent, time-series ratings of *The Nature Journey* is presented in Supplementary Figure S2. Scenes associated with awe and joy events showed significantly lower luminosity than control events (Figure 2F).

**Figure 2.**
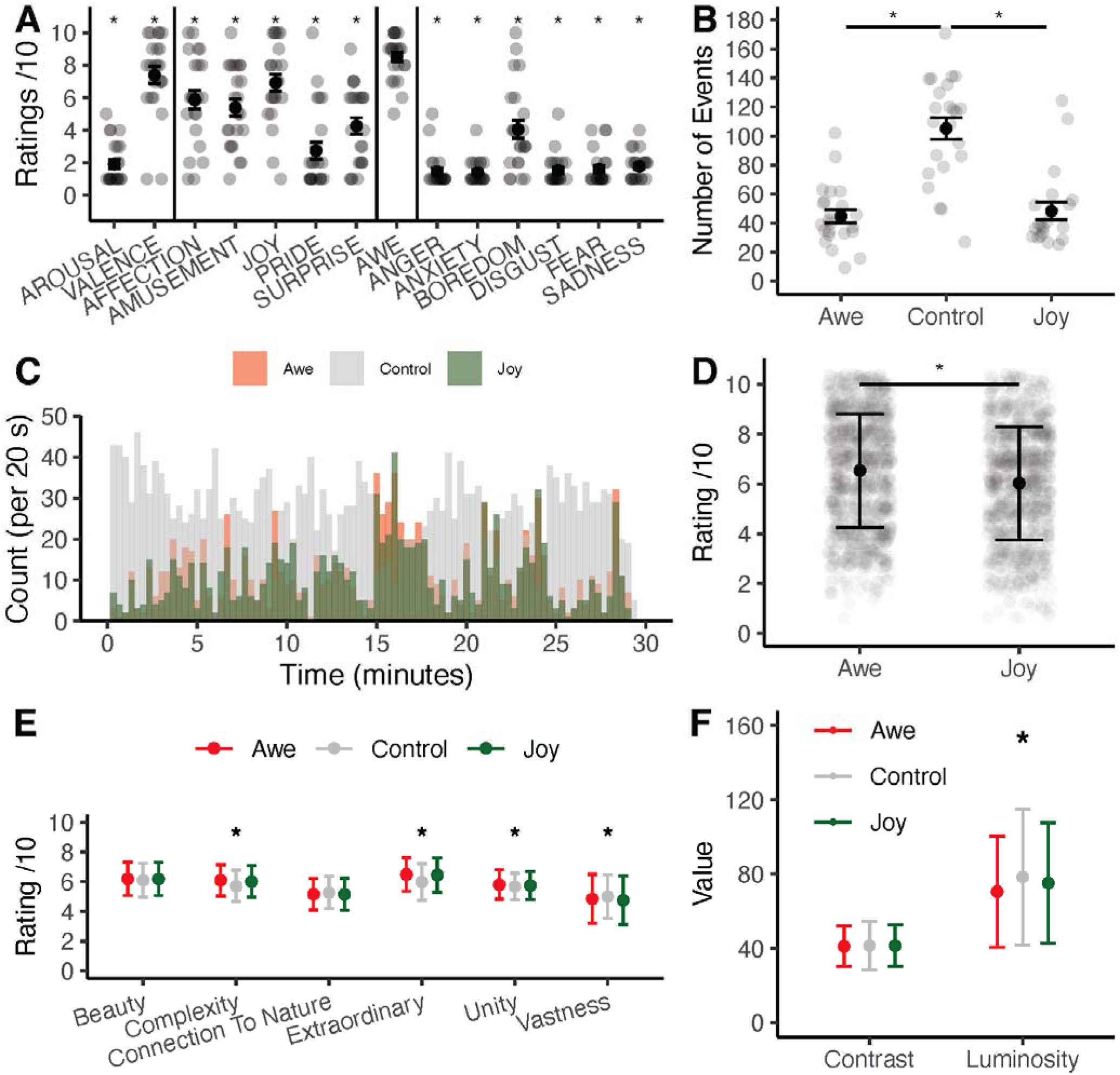
Emotional Characterization of The Nature Journey. (A) reports the ratings participants gave at the end of the first passive viewing of The Nature Journey. Asterisks denote significant difference from the awe rating. (B) reports the number of events for awe, joy, and control. (C) shows the distribution of awe, control, and joy events in 20 s bins across The Nature Journey. (D) compares the rating of awe and joy during the active viewing. (E) shows independent ratings of awe-relevant factors across scenes of The Nature Journey by six raters. The points and error bars reflect how the main study’s awe, control, and joy events differed by these independent ratings. (F) shows the difference in contrast and luminosity in the ±3 seconds surrounding awe, control, and joy ratings. Asterisks denote significant differences (p<0.05).

### 3.2. Neurophysiological Correlates of Awe

Analyses comparing EEG frequency bands between awe and control events revealed significantly decreased power in the theta (5 – 8 Hz, *d* = -0.54, *p =* 0.006, cluster-corrected) and alpha (8 – 13 Hz, *d* = -0.54*, p =* 0.007, cluster-corrected) spectral bands taking place approximatively -2.8 s to +0.6 s relative to awe events (Figure 3A). Topographical reconstructions revealed reduced theta and alpha power across mid-frontal and bilateral parietal regions (Figure 3B). Higher LZC under awe was observed when compared to control events in bilateral parietal electrode regions (*d* = 0.60, *p* = 0.022, cluster-corrected, Figure 3B).

**Figure 3.**
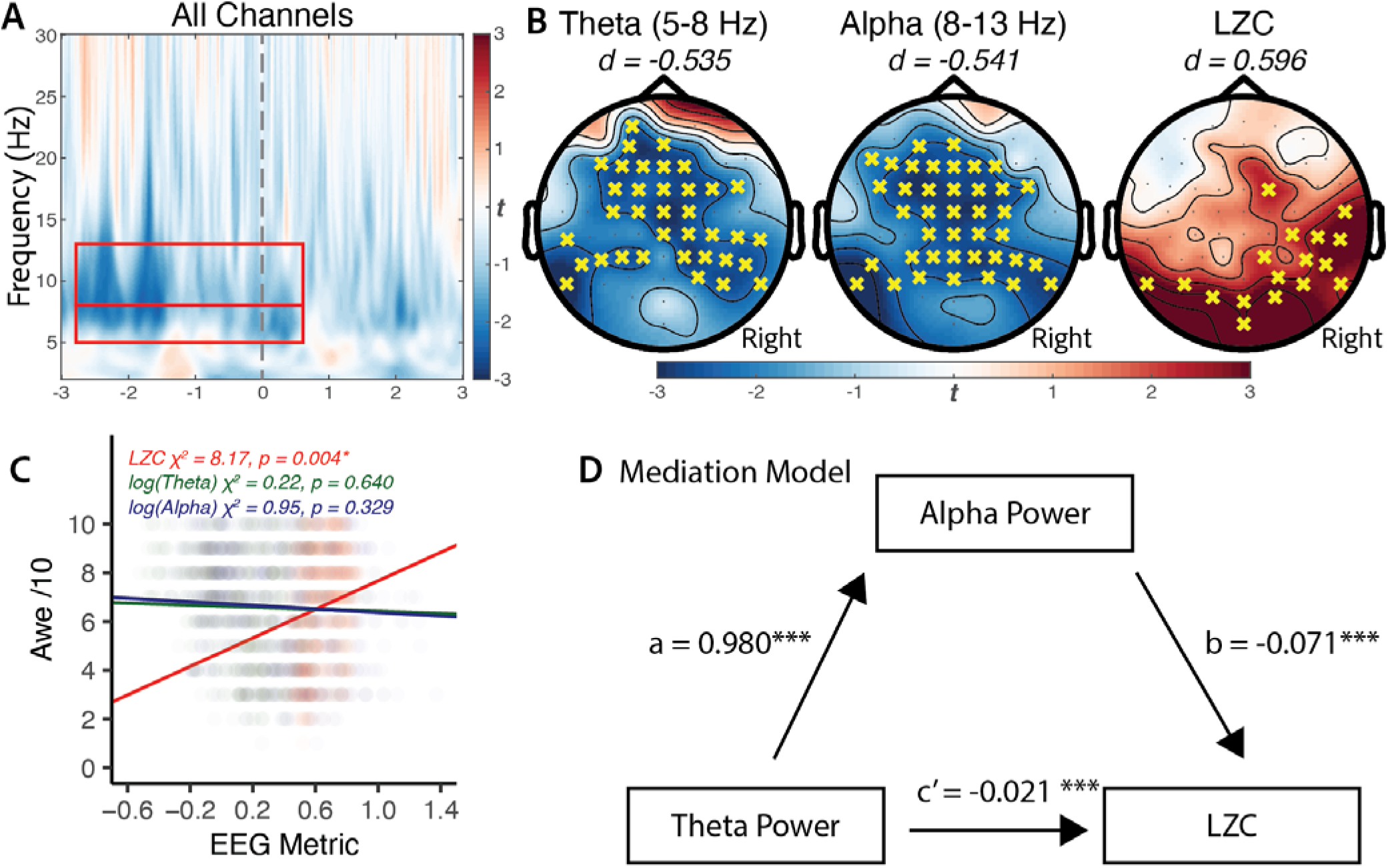
EEG-Based Neurophysiological Correlates of Awe. (A) Time frequency representation of paired statistical comparisons between averaged awe events against control events, averaged across all channels. The dashed grey vertical line at 0 s denotes the timing in respect to the occurrence of the awe or control event. The red rectangles (between -2.8 s and 0.6 s) show the theta (5 – 8 Hz) and alpha (8 – 13 Hz) frequency times and bands, which significantly differed between the awe and control conditions. The color bar depicts t-values, with blue indicating decreasing and red indicating increasing power. (B) Topographic statistical maps of theta, alpha, and Lempel Ziv Complexity (LZC) with yellow crosses denoting significant channels belonging to cluster-corrected comparisons between awe – control events. (C) shows that LZC significantly correlated with the intensity of awe ratings, but log(theta) and log(alpha) metrics did not. (D) shows mediation analysis where both decreases in theta and alpha power predict LZC increases, with alpha power partially mediating the relationship between theta power and LZC.

Further analysis assessed whether the intensity of the awe ratings corresponded to neurophysiological measures. Mixed effect models showed that LZC was positively associated with self-reported awe ratings (β = 2.93 ± 1.02, χ*^2^* = 8.17, *p* = 0.004, Figure 3C), but not theta (β = -0.21 ± 0.45, χ*^2^* = 0.22, *p* = 0.640, Figure 3C) nor alpha power (β = -0.36 ± 0.37, χ*^2^* = 0.95, *p* = 0.329, Figure 3C). However, as luminosity was lower in awe compared to control events (Figure 2F), a separate analysis controlling for luminosity as a fixed effect found that LZC remained significantly associated with awe (β = 2.62 ± 1.03, χ*^2^* = 6.49, *p* = 0.011, Supplementary Figure S1G).

A causal mediation analysis revealed that alpha power significantly mediated the relationship between theta power and LZC (Figure 3D). Theta power strongly predicted alpha power (β = 0.980, *t* = 33.50), and lower alpha power was associated with higher LZC after accounting for theta power (β = -0.071, t = -2.90). The average causal mediation effect (ACME) was significant (ACME = -0.070, 95% CI [-0.118, -0.023], *p* = 0.0046), whereas the average direct effect (ADE) of theta power on LZC was not significant after accounting for alpha power (ADE = -0.021, 95% CI [-0.078, 0.038], *p* = 0.475). The total effect of theta power on LZC remained significant (total effect = -0.091, 95% CI [-0.125, -0.056], *p* < 2e-16), with approximately 77% of the total effect mediated through alpha power.

Overall, these findings reveal a robust internal consistency of neurophysiological changes during awe, suggesting that the relationship between LZC increases and theta band decreases are mediated by decreases in the faster alpha band (Buzsáki & Vöröslakos, 2023). Crucially, the neurophysiological changes were not associated with contrast estimates from *The Nature Journey* (Supplementary Figure S1D,E,F). However, awe events were associated with lower luminosity (Figure 2F) and luminosity was negatively associated with LZC (Supplementary Figure 1A) – but even after accounting for luminosity effects, LZC was still associated with awe ratings (Supplementary Figure 1G). Overall, these findings suggest that awe-associated neurophysiological changes, are not the mere result of differences in low-level visual processes.

### 3.3. Source-Based Localization of Neurophysiological Correlates of Awe

Source-based localization analyses revealed that theta power decreases during awe events were primarily located within the dorsal, medial, and lateral parietal cortices as well as in the medial prefrontal and temporal cortices (Figure 4A). Alpha power decreases during awe were prominent in the visual, medial prefrontal, and temporal cortices (Figure 4B). LZC increases during awe were particularly present within the visual cortex, as well as in the dorsal, medial, and lateral parietal cortices (Figure 4C). Theta power decreases were primarily localized within the executive control network and the dorsal attention network (Figure 4D), while alpha power decreases were more widely distributed, being evident within the limbic, default mode, visual, executive control, and the dorsal attention networks (Figure 4E). Increases in LZC under awe were also widespread, particularly implicating the visual, executive control, and the dorsal attention networks (Figure 4F).

**Figure 4.**
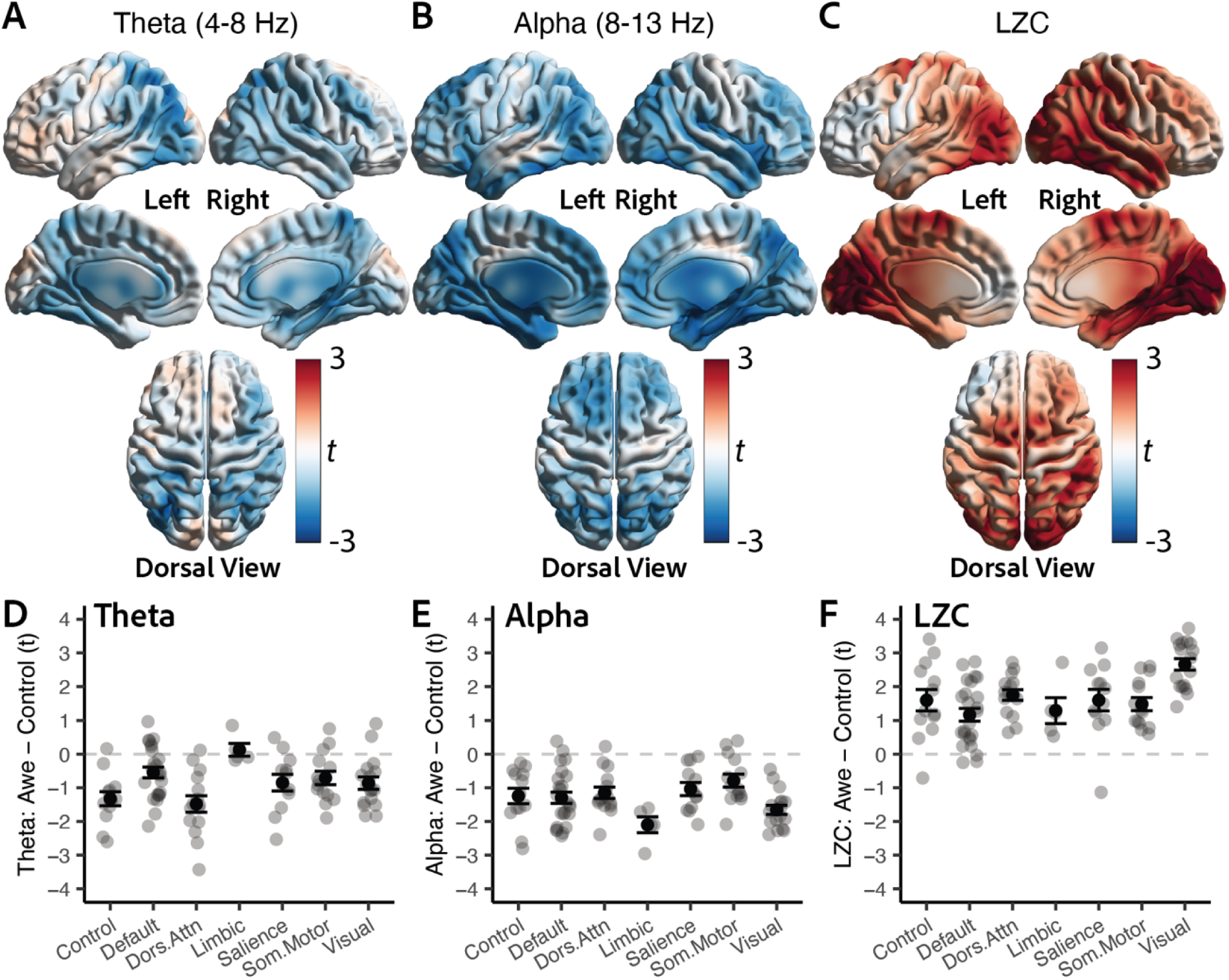
EEG Source-Based Neurophysiological Correlates of Awe. Source reconstruction performed using linearly constrained minimum variance beamformers and finite element models of individual T1-weighted MRIs to localize a grid of sources. The statistical comparison of awe-control values are shown for (A) theta power, (B) alpha power, and (C) Lempel Ziv Complexity (LZC). Changes in awe-control for (D) theta power, (E) alpha power, and (F) LZC are summarized for seven major intrinsic brain networks pertaining to the Schaefer atlas. Grey, translucent spheres indicate group-level region-of-interest (ROI) values, whereas black points denote mean of all networks’ ROIs, and error bars denote standard error. Control = executive control network; Default = default mode network; Dors.Attn = dorsal attention network; Limbic = limbic network; Salience = salience network; Som.Motor = somatomotor network; Visual = visual network.

### 3.4. Generalizability of Neurophysiological Correlates of Awe

Based on the primary findings presented in this study, the expected associations between awe ratings and EEG metrics were expected to be negative for theta, negative for alpha, and positive for LZC. The polarity of these associations was only partially generalizable across the different studies (Figure 5A). For theta, negative associations with awe ratings were observed in the Hu, 2017 dataset, but not observed in the present study (Figure 3C and Figure 5) whereas positive awe rating associations were observed in the Emotion Rating Study (ERS) and DMT datasets (Figure 5). For alpha, negative alpha associations with awe were observed in the present study and the Hu, 2017 dataset, but a mixed positive and negative awe topographic association was observed in the ERS and DMT datasets (Figure 5). For LZC, the present study, ERS, and DMT studies showed positive occipital associations with awe, whereas the Hu, 2017 dataset showed a general negative association with awe (Figure 5).

**Figure 5.**
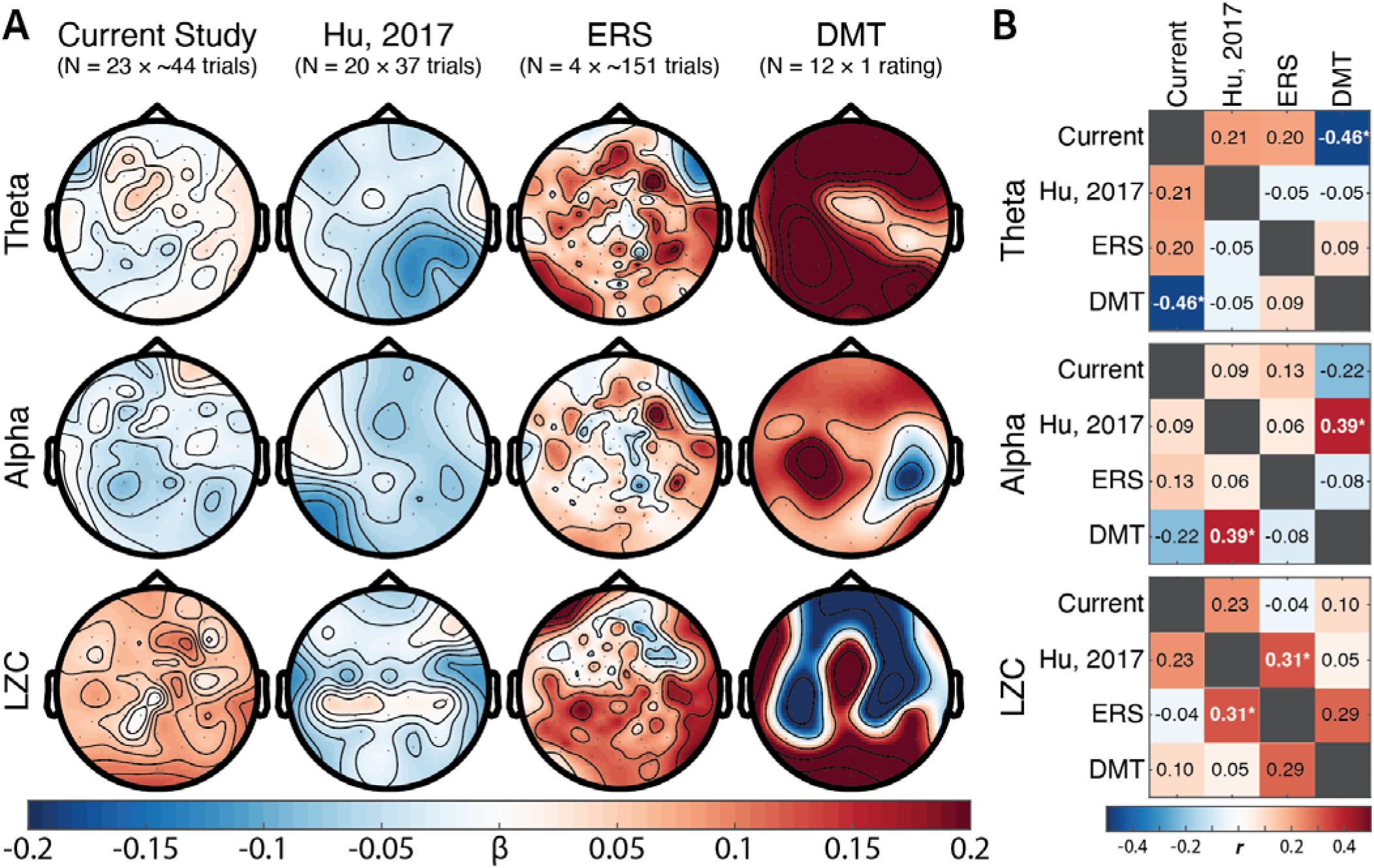
Generalizability of Neurophysiological Correlates of Awe. (A) Standardized beta linear model analysis of awe ratings predicted by theta, alpha, and Lempel Ziv Complexity (LZC) per channel. Four studies are included: (i) the current study (N = 23, ∼44 ratings each), (ii) published data from Hu, 2017 (N = 20, 37 ratings each), (iii) unpublished data from an Emotion Rating Study (ERS, N = 4, ∼151 ratings each), and (iv) published data investigating the effects of N,N-dimethyltryptamine (DMT) from Timmermann et al., (2019), N = 12 with 1 awe rating each. (B) Heatmaps show the pairwise spatial similarity correlation between the four studies across the three EEG metrics with asterisks denoting permutation-based significant spatial correlations between topographies (p<0.05).

A supplementary pilot analysis (N = 1) assessed the experience of goosebumps – a phenomenon often accompanying awe (Benedek & Kaernbach, 2011; Maruskin et al., 2012), during a 40 min audiovisual stimulus (Supplementary Figure S3). While these results were only in one participant, reduced theta, reduced alpha, and increased occipital LZC was observed in goosebump events compared to control. However, as these events only mark the events of goosebumps, rather than the intensity of any awe rating, comparable topographies were not included in Figure 5.

### 3.5. Uniqueness of LZC as a Neurophysiological Correlate of Awe Compared to Joy

As joy was the second-highest rated emotion after the first viewing (Figure 2A), joy was selected as a second emotion to rate in a secondary experiment, of which 20 of 23 participants completed. Joy events also showed decreased theta power (though not cluster significant), decreased alpha power (Cohen’s *d* = -0.52, *p =* 0.03, cluster-corrected), and increased LZC (*d =* 0.57, *p* = 0.03, cluster-corrected) compared to control events. Using the LZC values from channels belonging to the significant cluster, awe ratings exhibited a positive association with LZC (β = 4.7, χ*^2^* = 5.21, *p* = 0.022), but interaction effects, testing for the difference between joy and awe slopes, did not show significantly different slopes (joy slope β*1* = 4.8, interaction χ*^2^*= 1.15, *p* = 0.283) (Figure 6C). While awe showed larger theta decreases, alpha decreases, and LZC increases, these comparisons did not significantly differ between awe or joy events.

**Figure 6.**
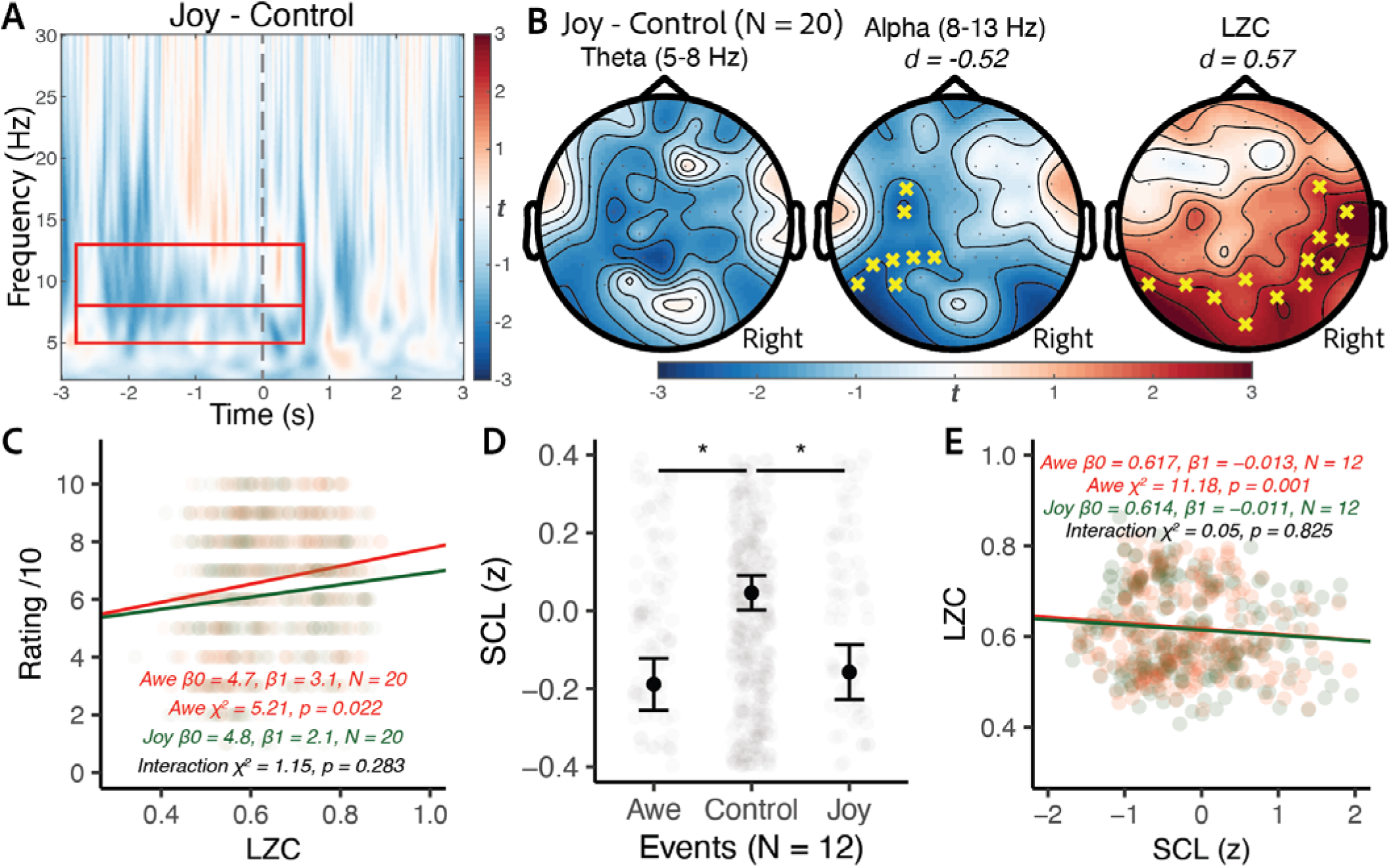
Uniqueness of the Neurophysiological Correlates of Awe. (A) EEG time frequency representation of the joy minus control events. Red rectangles denote the same time-frequency areas which showed maximal awe effects. (B) shows topography of theta, alpha, and Lempel Ziv Complexity (LZC) measures relative to control events. Yellow crosses denote channels belonging to statistically significant clusters. Cohen’s d effect sizes are shown for the significant clusters. (C) shows that awe is significantly predicted by LZC measures but the joy slope does not significantly differ from the awe slope. (D) shows skin conductance level (SCL) is lower in awe and joy events compared to control events. (E) shows that LZC is negatively associated with SCL in awe events, but that the LZC/SCL slopes do not significantly differ.

### 3.6. LZC under Awe Events Predicted by Reduced Sympathetic Activity

Both awe events (β*1 = -0.23 ± 0.06 (s.e.), p = 0.0008)* and joy events (β*1 = -0.20 ± 0.07 (s.e.), p = 0.0009*) showed lower skin conductance compared to control (Figure 6E). Since LZC was positively associated with awe, and awe events had lower SCL compared to control, we anticipated a negative relationship between LZC and SCL. This negative relationship between LZC and SCL was observed (β*1 = -0.013 ± 0.004 (s.e.), p = 0.0008*). However, to ascertain whether SCL exhibited a differing relationship with joy, interaction analyses showed that the relationship between LZC and SCL was not significantly different in awe events compared to joy events (χ^2^ = 0.05, *p =* 0.825). To ensure that luminosity did not confound the results, SCL still remained significantly associated with LZC even when controlling for luminosity as a fixed effect (β*1 = -0.012 ± 0.004 (s.e.), p = 0.0015*, Supplementary Figure S1H).

### 3.7. Awe and LZC: Trait Characteristics or Transient States

The mean awe intensity self-reported by participants negatively correlated with rumination and loneliness (Supplementary Figure S4B,C). This suggests that self-reported awe experience reflects to some degree an individual trait. However, when correlating the mean LZC during awe events, LZC did not correlate with reflection, rumination or loneliness (Supplementary Figure S4D,E,F) – suggesting that neurophysiological changes associated with the awe experience reflect a transient state rather than an individual personality trait.

## 4. Discussion

Awe, and the feelings of wonder, vastness and unity that often accompany this emotion, can catalyze transformative experiences, prompting people to re-evaluate their view of themselves and the world, with potential positive effects on well-being and prosocial behavior (Keltner & Haidt, 2003; Monroy & Keltner, 2022; Shiota, 2021; Shiota et al., 2007). Yet, the neural correlates of awe have so far remained elusive (X. Hu et al., 2017; Pasquini et al., 2023; van Elk et al., 2019). The present findings suggest that our nature-inspired audiovisual stimulus – *The Nature Journey* – primarily elicits awe along with various other positive emotions (Figure 2). Participant-rated awe events, in comparison to randomly selected control events, exhibit reduced theta power, reduced alpha power, but increased LZC – with only LZC showing a significant association with the intensity of awe ratings (Figure 3). Awe events exhibit lower skin conductance than control events, and LZC also exhibits an expected negative association with SCL (Figure 6). When comparing the LZC-awe association topographies across studies, occipital LZC is positively associated with awe ratings (Figure 5). However, secondary analysis to explore whether LZC increases were a unique signature of awe showed that joy ratings and joy events also show similar neurophysiological changes. Overall, these findings suggest that experiences of awe may have underlying neurophysiological markers that, while potentially not entirely unique to awe, are characterized by increases in complexity of activity in posterior brain areas.

### 4.1. *A*we and Other Positive Emotions are Induced through Naturalistic Stimuli

In this first characterization of *The Nature Journey*, awe emerged as the predominant elicited emotion. Although participants also reported substantial feelings of joy (Figure 2A,D), awe was consistently rated as the most intense emotion, indicating that the intervention selectively elicited awe rather than broadly increasing positive affect. Notably, *The Nature Journey* elicited several experiential qualities commonly associated with awe, including perceptions of complexity, extraordinariness, and feelings of unity, despite comparatively lower ratings of vastness (Figure 2E). Contemporary accounts increasingly emphasize awe as a self-transcendent emotion characterized by diminished self-focus and a heightened sense of connection to others and the broader world (Stellar et al., 2017). Viewed through this lens, the present findings suggest that *The Nature Journey* may evoke awe through pathways other than perceptions of overwhelming physical scale. Rather, engagement with complex and extraordinary natural phenomena may promote a shift away from habitual self-referential processing toward a broader and more interconnected perspective. Consistent with this interpretation, higher mean awe ratings were associated with lower loneliness and reduced self-directed rumination (Supplementary Figure S4). These findings align with theoretical accounts proposing that awe’s psychological benefits are mediated through reductions in self-focus and increases in social connectedness (Monroy & Keltner, 2022). Together, our results suggest that *The Nature Journey* evokes a form of awe rooted less in perceived vastness and more in self-transcendent experiences of unity, interconnectedness, and engagement with the complexity of the natural world.

### 4.2. Skin Conductance and EEG Theta and Alpha Changes Under Awe Reflect Low Arousal and Positive Valence

The present EEG findings align with past research characterizing awe as a low arousal and positively valenced emotion. *The Nature* Journey was characterized by ratings of overall low arousal and high valence (Figure 2A) and participants exhibited reduced theta power (Figure 3B) and reduced alpha power (Figure 3B). Conversely, theta power is enhanced by high arousal (Aftanas et al., 2002, 2004; Balconi & Lucchiari, 2006; Balconi & Pozzoli, 2009) and negative valence (Aftanas et al., 2001). Although alpha power decreases have been associated with high arousal (Bonnet & Arand, 2001; Martin et al., 2019), awe has been understudied, with one single previous study assessing the neurophysiological correlates of 10 positive emotions by correlating the individual topographies induced by short video-clips with the self-reported strength of the experienced emotions (X. Hu et al., 2017). This study revealed that higher subjective intensities of reported awe correlated with reductions of all major EEG frequency-bands, including decreases in the theta and alpha power bands, in line with our study (Figure 3). Several cognitive processes can induce frontal-midline as well as posterior theta and alpha desynchronization, including semantic and memory processes (Herweg et al., 2020; Klimesch et al., 1994, 1997), as well as increased alertness and attentional demands (Klimesch et al., 1998; Palva & Palva, 2007; Peylo et al., 2021; Woodman et al., 2022). Theta and alpha power decreases in our study did not correlate with self-reported awe strength (Figure 3C) suggesting that these neurophysiological changes may be induced by a variety of cognitive processes not specific to the experience of awe. This is consistent with recent reviews arguing that alpha desynchronization indexes more general emotional processes and engagement of motivational systems (Codispoti et al., 2023). Aligned with this narrative, increased theta-band connectivity between the default mode and frontoparietal control networks has been linked to internally directed attention (Kam et al., 2019). Given the overlap between the frontoparietal control network and regions of the control and dorsal attention networks (Dixon et al., 2018), the reduction in theta power observed across both control and dorsal attention networks (Figure 4D) may reflect a shift away from internally focused cognition and toward heightened engagement with the external environment. Concurrently, parietal alpha suppression has been associated with the processing of conflict and the allocation of attentional resources to salient information (Cohen & Ridderinkhof, 2013). The reduction in alpha power observed in our study (Figure 4E) may therefore indicate increased processing of information that challenges existing expectations. Together, these findings suggest a neurophysiological state that may facilitate accommodation, a core component of awe in which prior mental schemas are revised in response to experiences perceived as vast or extraordinary.

The present findings pertaining to skin conductance corroborate the low arousal nature of awe. Previous studies assessing the effect of awe on autonomic physiology have consistently reported reduced sympathetic activity (Chirico et al., 2017; Monroy & Keltner, 2022; Shiota et al., 2011) while participants experience awe, typically assessed through decreases in SCL – a purely sympathetically driven signal (Clark et al., 2018; Hernes et al., 2002). Our findings showing that SCL was lower for awe events (Figure 6D) further corroborate previous reports of reduced sympathetic activity under awe (Behnke et al., 2022).

### 4.3. LZC Increase as a Potential Marker for Awe

We propose LZC increase as a potential neurophysiological marker for awe. LZC was observed to be higher in awe compared to control events (Figure 3B). LZC was associated with the intensity of awe ratings (Figure 3C), as well as negatively associated with SCL (Figure 6E) – which has been shown to be lower in awe (Kreibig, 2010; Shiota et al., 2011). LZC is increasingly used in the field of cognitive and consciousness neuroscience to measure the unpredictability of brain activity (Schartner, Carhart-Harris, et al., 2017; Schartner, Pigorini, et al., 2017) and may reflect greater information processing capacity or more diverse neural populations (Höhn et al., 2024). Lower LZC has been reported in patients with disorders of consciousness, Alzheimer’s disease dementia, as well as under unconscious states like anesthesia or deep sleep (Aamodt et al., 2023; Abásolo et al., 2006; Boncompte et al., 2021; Hudetz et al., 2016; Liu et al., 2023). Conversely, higher LZC has been found under meditative states (D’Andrea et al., 2024; Martínez Vivot et al., 2020) and when under the effects of psychedelics (Schartner, Carhart-Harris, et al., 2017; Timmermann et al., 2023), both states of expanded conscious experiences – that are also often associated with feelings of awe (Preston & Shin, 2017; Tsaur et al., 2024). In particular, the similarity of LZC topographies found under awe in our study and DMT supports the idea of a shared neurophysiology (Goldy et al., 2024; Hendricks, 2018; Yaden et al., 2024). This biological convergence might explain – and potentially validate – why psychedelics, meditation, and awe can all have positive impacts on mental health and well-being (Bogenschutz et al., 2022; R. Carhart-Harris et al., 2021; Falchi-Carvalho et al., 2025; Goodwin et al., 2022; Griffiths et al., 2016; Raison et al., 2023).

Awe-related increases in LZC may encapsulate the rich sensory experience elicited by *The Nature Journey*. Source localization revealed LZC increases within posterior cortical regions encompassing the visual, dorsoparietal, and medial parietal cortices, overlapping with large-scale brain systems including the visual, executive control, dorsal attention, and default mode networks (Buckner & DiNicola, 2019; Uddin et al., 2019; Vossel et al., 2014). Increased activity within the visual network is expected given the rich audiovisual nature of the stimulus (Chang et al., 2015; Nishimoto et al., 2011). Because LZC has been proposed to index the richness and diversity of conscious experience (R. L. Carhart-Harris, 2018), elevated LZC within visual regions may reflect the complexity and extraordinariness of the naturalistic scenes presented in *The Nature Journey* (Figure 2B).

Awe-related increases in LZC may potentially also reflect the cognitive processes underlying awe-related changes in self-perspective. The present study found LZC increases in the posterior cingulate and retrosplenial cortices (Figure 4C), regions implicated in self-referential functions (Gallagher, 2000; Lyu et al., 2023; Northoff & Bermpohl, 2004). Within the “relaxed beliefs under psychedelics” (REBUS) framework, increases in neural complexity are proposed to reflect a loosening of high-level priors and greater sensitivity to bottom-up sensory information (R. L. Carhart-Harris & Friston, 2019). As such, the awe-related LZC increases in these self-referential regions may promote temporary reorganizations of self-related beliefs – implicated in the diminished “sense of self” induced by awe (Bai et al., 2017; Monroy & Keltner, 2022; Piff et al., 2015). Indeed, prior task-based fMRI investigations found awe-related decreases in default mode network activity (van Elk et al., 2019). Altered dynamics within these self-referential regions may therefore relate to the reduced self-focus and “small self” experiences frequently associated with awe (Bai et al., 2017; Monroy & Keltner, 2022; Piff et al., 2015). Increased LZC was also observed within dorsoparietal regions involved in spatial attention, self-location, and the representation of one’s relationship to the surrounding environment (Corbetta & Shulman, 2002; Husain & Nachev, 2007; van Elk et al., 2016). This interpretation aligns with theoretical accounts describing awe as a self-transcendent state characterized by reduced self-focus and greater connectedness to the broader world (Bai et al., 2017; Monroy & Keltner, 2022).

### 4.4. Generalizability of Findings and Mixed Emotions

A central debate in the field of affective science is whether a physiological pattern associated with a specific emotion can be generalized across experimental procedures and study populations (Kragel & LaBar, 2016; Kreibig, 2010; Lindquist et al., 2012; Siegel et al., 2018). Functionalist models of emotions propose, on the one hand, that emotions are rooted in evolutionary biology, viewing emotions as brief and dynamic states that involve distinct experiences, expressive behaviors, patterns of thought, and physiological changes which evolved as adaptive responses with specific functions for survival and social interaction (Adolphs, 2017; Adolphs & Andler, 2018; Ekman, 1992; Gross, 2015; Lench et al., 2011; Scherer, 2005). Constructivism, on the other hand, emphasizes the role of social and cultural context in shaping emotional experiences, suggesting that emotions are not innate but rather constructed through social learning and interpretation (Adolphs et al., 2019; Barrett, 2017; Jackson et al., 2019; Lindquist et al., 2022; J. A. Russell & Barrett, 1999). This debate is still ongoing, primarily driven by the limited generalizability of physiological signatures of emotions, as demonstrated by large metanalyses reporting incongruent findings when systematically assessing for the presence of characteristic physiological patterns of emotions across studies (Barrett, 2017; Lindquist et al., 2012; Siegel et al., 2018). Our finding that occipital LZC increased during awe across multiple datasets – spanning younger and older adults with varying number of trials, different audiovisual stimuli, and pharmacologically induced awe – provides evidence for a cross-sample neurophysiological marker.

Nevertheless, important differences were found across datasets, in particular when relating the topographies of neurophysiological changes found in our study of US-based participants to those derived from a Chinese dataset (X. Hu et al., 2017). While we observed some degree of cross-cultural convergence in the theta and alpha power topographies, we also found notable divergence in LZC in the Chinese sample. These differences could be arising from methodological differences, such as the use of shorter videoclips, or reflect the influence of cultural backgrounds and social factors in the experience of emotions (Barrett, 2017; Lindquist et al., 2012; Siegel et al., 2018). For example, Chinese participants typically report higher levels of fear when experiencing awe (Mandarin: 敬畏, Jìngwèi) compared to US participants (Stellar et al., 2024). Prior work suggests that individuals from an East Asian cultural context are more likely to experience awe as a mixed emotion, encompassing both positive and negative elements (Fang & Kawakami, 2025). More broadly, there is an increased recognition that awe, as well as other emotions, may be experienced in unison with other emotional states (Stellar et al., 2024). This complex emotional experience, where mixed emotions at times of contrasting valence may be experienced at varying strengths (Berrios et al., 2015; Kreibig et al., 2013; Kreibig & Gross, 2017; Larsen & McGraw, 2014), has been shown to be an important moderator of mood disorders and mental well-being (Berrios et al., 2018; Braniecka et al., 2014; Tan et al., 2022). Awe has been shown to be an emotion that can take various forms, varying from positive awe, as induced by *The Nature Journey* used in our study, to threat-based awe, as experienced for example in the presence of a catastrophic event (Cowen & Keltner, 2021; Gordon et al., 2017; Monroy & Keltner, 2022). Although our study lays the groundwork for exploring the neurophysiological basis of nature-inspired awe, future studies are needed to deepen our understanding of the neurophysiological signatures induced by distinct states of awe.

### 4.5. Study Limitations – Uniqueness of Signal and Limited Control Event Comparison

Caution should be applied interpreting the present findings due to the apparent lack of specificity for the neurophysiological measures of awe. To assess the uniqueness of the observed effects, a secondary experiment was conducted in which twenty of the twenty-three participants additionally rated moments of joy on a separate visit. Remarkably, joy ratings exhibited patterns similar to those observed for awe across multiple neurophysiological measures, although the corresponding effect sizes were generally smaller (Figure 2C, Figure 6A–C). While awe was the predominant emotion elicited by *The Nature Journey*, these findings suggest that the observed changes in LZC, spectral power, and autonomic arousal may not be uniquely specific to awe. Rather, they may reflect broader processes shared across positive emotional experiences, such as attentional engagement, experiential richness, emotional intensity, or self-transcendence. Consequently, the overlap observed in the present study may indicate either that the measured neurophysiological changes reflect processes common to multiple positive emotions or that the participants were experiencing blended awe-joy states in response to the stimulus. Future studies designed to more clearly dissociate awe from related positive emotions will be necessary to determine the extent to which these neurophysiological signatures are specific to awe.

Furthermore, the study design is limited due to the variability in defining control events. As the control events are unspecific periods of the movie where an individual participant did not specify an awe event, the range of possible subjective experiences and thoughts is highly variable. The original intent was to retain a context-relevant control by using neurophysiological recordings of participants watching the same movie as opposed to resting-state eyes open recordings. Further, as the number of control events far outweigh number of awe events and were randomly selected for each participant (Figure 2B), this procedure would ideally retain a stable, representative, sample of control events. Future studies may benefit from experimental designs where participants actively note moments of interest when they feel something during the first viewing and retroactively return to these small snippets afterwards to rate how they felt.

### 4.6. Conclusion

Taken together, these findings identify a low arousal, positively valenced, neurophysiological signature of awe, centered on increased neural entropy in posterior cortical networks and linked to both the subjective experience and autonomic physiology. We interpret our data as implying a role for increased information processing in higher-level cortical systems in this phenomenology of awe, consistent with prior work assessing the neural basis of a diminished “sense of self”.

## Data availability

Data will be made available upon publication at an openly accessible repository.

## Code availability

The code used to analyze is uploaded in https://github.com/josephccchen/dynamo.

## Acknowledgements

We thank the participants of the study for their invaluable contribution to research. We thank the artists Louie Schwartzberg and East Forest for creating the audiovisual recordings used to generate the primary data analyzed in this study.

