## Supplement for "Complexity as a Potential Neurophysiological Correlate of Awe"

### Supplementary Materials

Supplementary Table 1. Sample demographics and characterization. 23 participants were recruited in this study. Mean scores and standard deviation for demographical information and standardized neuropsychological tests used as criteria to assess healthy cognitive performance. Each participant scored within -1.5 SDs of their age-matched normative value for minimum performance.

| Demographics | |
| --- | --- |
| Age in years | 70.4 ± 6.4 |
| Sex | 11 M, 12 F |
| Education in years | 17.8 ± 1.9 |
| Income in US $  Below $100,000  $100,000 - $200,000  $200,001 - $300,000  $300,001 - $400,000 | 6  14  2  1 |
| Ethnicity  White  Asian | 19  4 |
| Remote Neurological Assessment (z) | |
| CVLT Immediate Recall | 0.55 ± 0.96 |
| CVLT Short | 1.16 ± 0.76 |
| Digit Span Backwards | -0.19 ± 0.73 |
| Lexical Fluency | -0.02 ± 0.82 |
| Semantic Fluency | -0.35 ± 0.82 |
| Trails | -0.09 ± 1.2 |
| CVLT Long | 0.91 ± 0.84 |
| CVLT Cued | 0.62 ± 1.38 |
| Neuropsychological Surveys | |
| UCLA Loneliness 20 | 7.1 ± 6.6 |
| RRQ-Reflection | 33.8 ± 7.9 |
| RRQ-Rumination | 29.2 ± 8.8 |

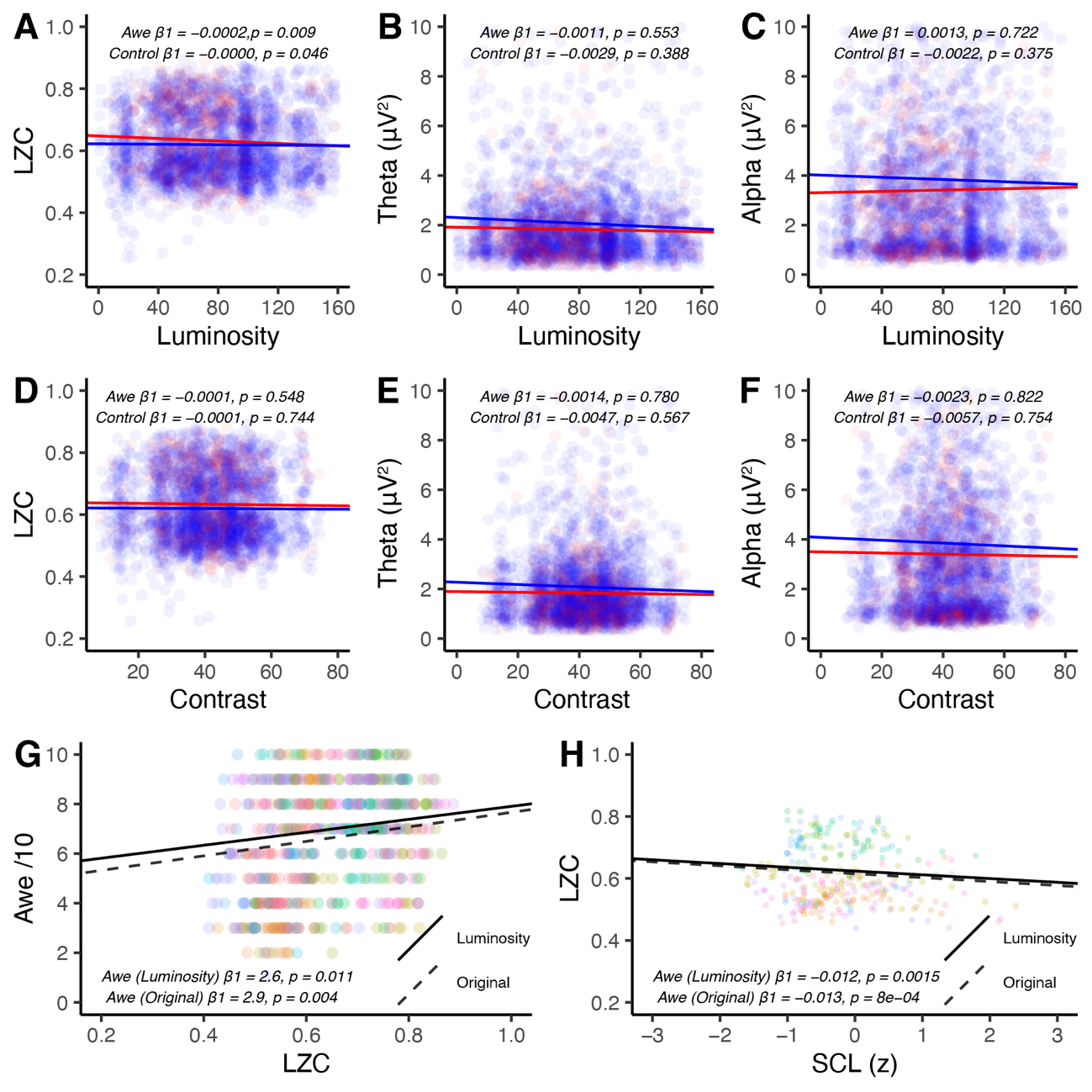

Figure S1. Electroencephalographic (EEG) Measures’ Association with Visual Features. Linear mixed effect models were performed associating EEG metrics to luminosity and contrast measures. In (A) – (F), EEG metric ~ Feature*Type + (1|ID) was performed where Feature was either luminosity or contrast and Type represented either awe or control events. The p-values denote the significance of the awe events’ slope, and the interaction to see whether the control slope differed from the awe slope. (A) shows Lempel Ziv Complexity (LZC) was associated with luminosity in awe events, but the LZC slope in control events significantly differed from the awe slope (p = 0.046). (B) – (F) do not show significant associations of the visual features with other EEG metrics. (G) shows that the awe rating retained a significant relationship with LZC measure even when accounting for luminosity. (H) shows that the relationship between LZC and skin conductance level (SCL) retained significance even when accounting for luminosity.

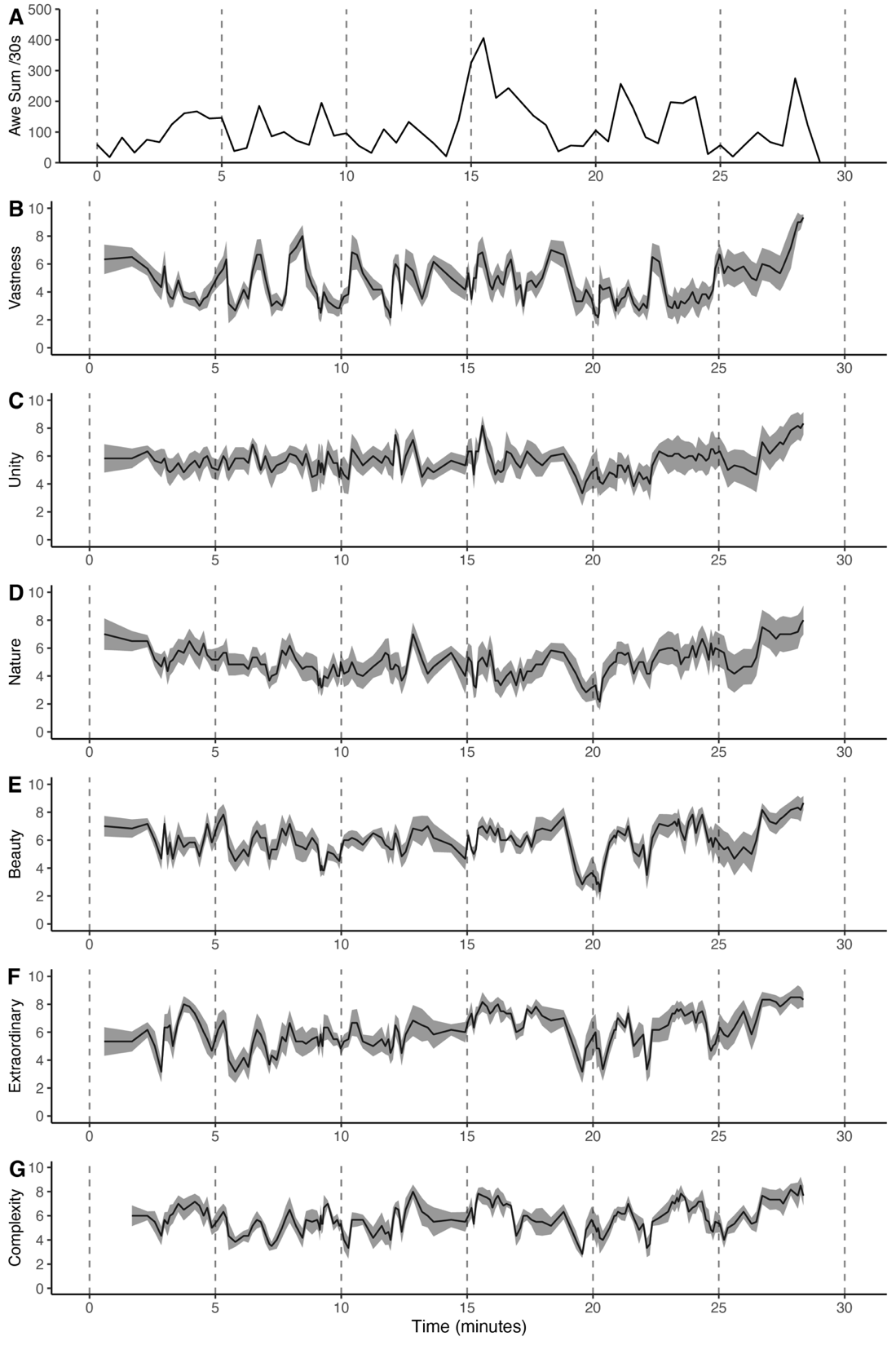

Figure S2. Independent ratings of movie scenes. (A) Ratings of awe for 30 sec epochs summed across all study participants. Six independent raters rated each scene of The Nature Journey for feelings of (B) vastness, (C) unity, (D) connection to nature, (E) beauty, (F) extraordinary, and (G) complexity. Time series for these constructs across the duration of the entire movie are plotted, with black denoting the group mean and gray the standard deviation.

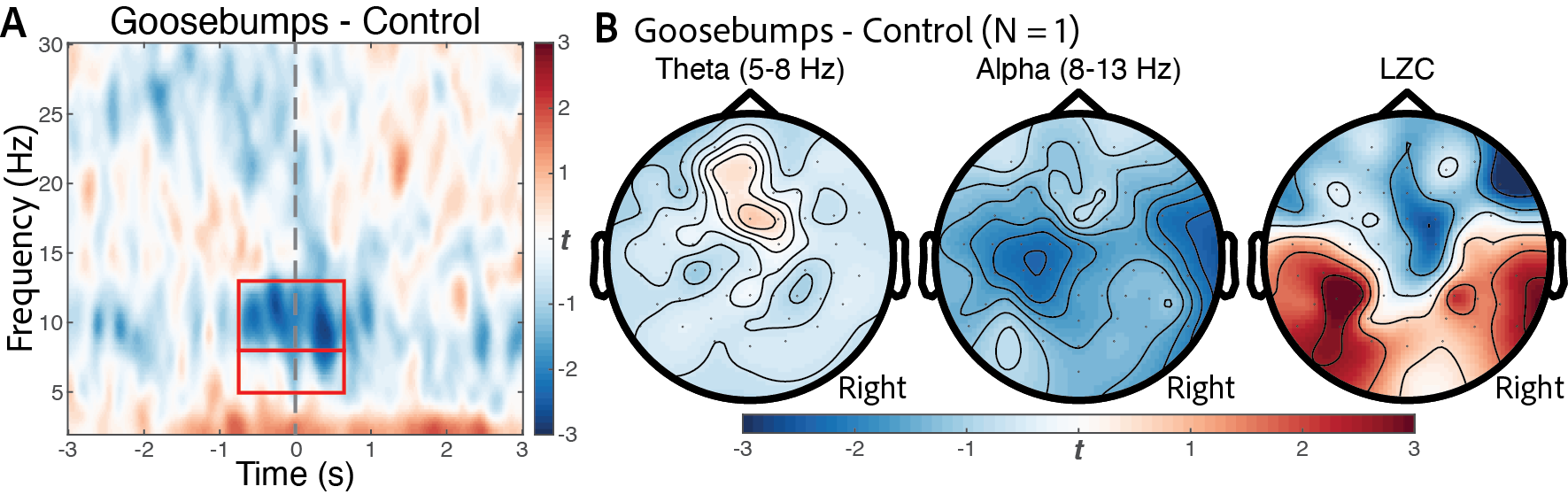

Figure S3 – Electroencephalography (EEG) Measure of Goosebumps. Self-reported goosebump events are compared to pseudorandomly generated control events pertaining to timepoints not characterized by goosebumps. (A) Statistical time frequency representation of goosebumps compared to control events, averaged across all 64 channels. (B) shows the theta, alpha, and Lempel Ziv Complexity (LZC) measures derived from the -0.5 to +0.5 s around each goosebump event.

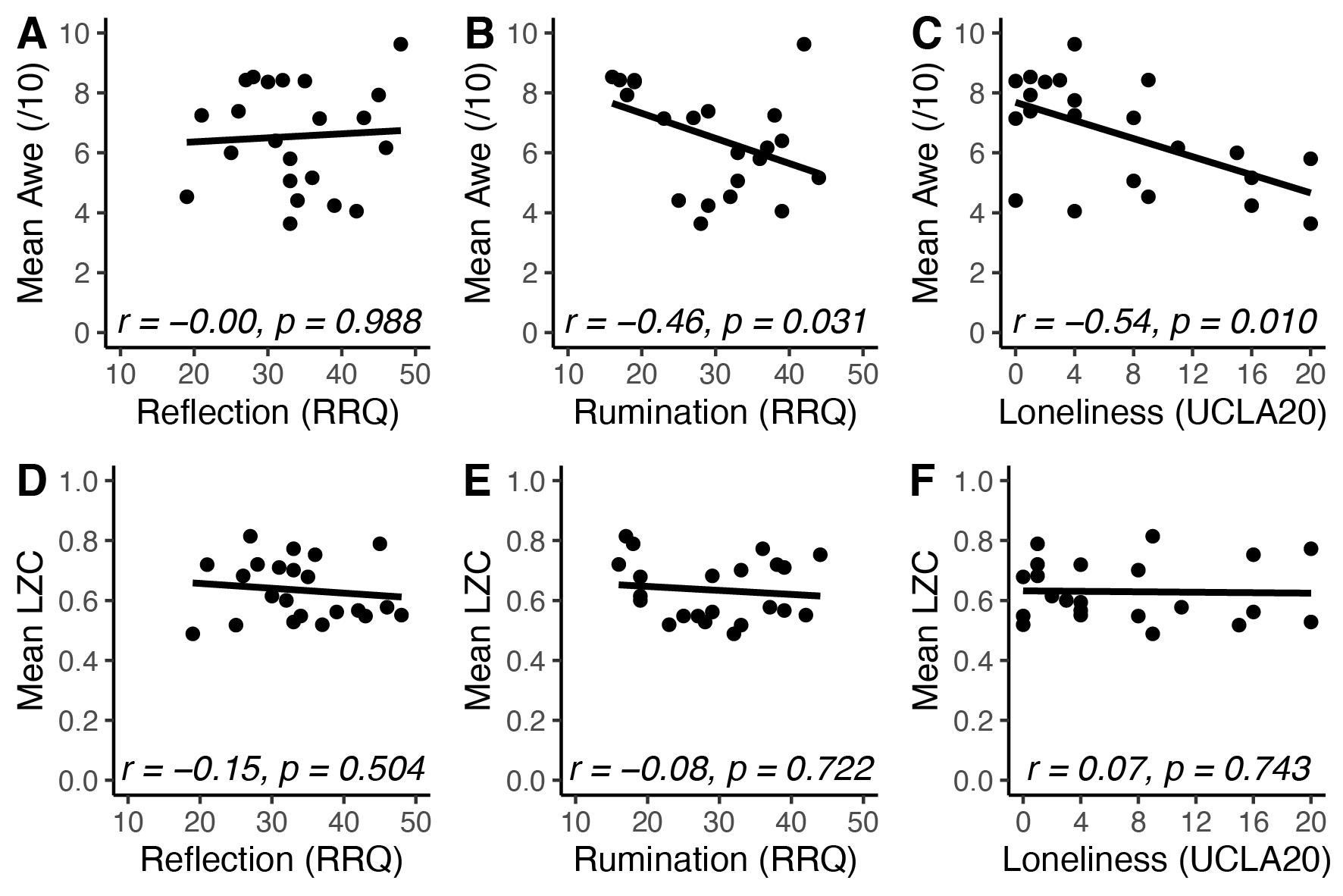

Figure S4 – Self-referential and loneliness scores. (A) Scores for reflection from the RRQ were not significantly associated with awe intensity averaged across awe events. (B) Scores for rumination from the RRQ were significantly associated with awe intensity averaged across awe events. (C) Scores for loneliness from the UCLA loneliness scale 20-items were significantly associated with awe intensity averaged across awe events. LZC complexity averaged across awe events was not significantly associated with reflection (D), rumination (E), or loneliness (F). RRQ: Rumination Reflection Questionnaire.
